# Cell cycle progression mechanisms: slower cyclin-D/CDK4 activation and faster cyclin-E/CDK2

**DOI:** 10.1101/2023.08.16.553605

**Authors:** Wengang Zhang, Yonglan Liu, Hyunbum Jang, Ruth Nussinov

## Abstract

Dysregulation of cyclin-dependent kinases (CDKs) impacts cell proliferation, driving cancer. Here, we ask why the cyclin-D/CDK4 complex governs cell cycle progression through the longer G1 phase, whereas cyclin-E/CDK2 regulates the short G1/S phase transition. We consider the experimentally established high-level bursting of cyclin-E, and sustained duration of elevated cyclin-D expression in the cell, available experimental cellular and structural data, and comprehensive explicit solvent molecular dynamics simulations to provide the mechanistic foundation of the distinct activation scenarios of cyclin-D/CDK4 and cyclin-E/CDK2 in the G1 phase and G1/S transition of the cell cycle, respectively. These lead us to propose slower activation of cyclin-D/CDK4 and rapid activation of cyclin-E/CDK2. Importantly, we determine the mechanisms through which this occurs, offering innovative CDK4 drug design considerations. Our insightful mechanistic work addresses the compelling cell cycle regulation question and illuminates the distinct activation speeds in the G1 versus G1/S phases, which are crucial for cell function.

**Statement:** Our work provides an unprecedented mechanistic understanding of the distinct activation scenarios of cyclin-D/CDK4 and cyclin-E/CDK2 in cell cycle regulation, underpinning the slower activation of cyclin-D/CDK4 in the more extended G1 phase and the rapid activation of cyclin-E/CDK2 in the brief G1/S transition. Our findings address a long-standing question in cell cycle biology and suggest the design of targeted CDK4 inhibitors.

## 1 | INTRODUCTION

Cyclin-dependent kinases (CDKs) are a family of serine/threonine kinases essential for cell cycle regulation.(Malumbres et al. 2009; Morgan 2007; Patterson et al. 2021) Among them, CDK2 and CDK4 are crucial for the initiation of DNA replication and progression from the G1 to the S phase. The kinases are activated by binding to their respective regulatory cyclin partners; cyclin-A and cyclin-E to CDK2 and cyclin-D to CDK4. Cyclins are synthesized and degraded in a cyclical fashion throughout the cell cycle and cyclin/CDK complexes are further regulated by phosphorylation and dephosphorylation events. Aberrant expressions of cyclin-D, cyclin-E, and cyclin-A are common in cancer cells.(Gallo et al. 2022) Cyclin/CDK complexes undergo specific changes in their activity during different stages of the cell cycle, ensuring proper cell cycle progression. Dysregulation of CDK activity is frequently observed in cancer cells and plays critical roles in the development and progression of cancer and neurodegenerative disorders.(Lukasik et al. 2021; Malhotra et al. 2021; Malumbres and Barbacid 2009; Matthews et al. 2022; Otto and Sicinski 2017; Wander et al. 2022) CDK2 and CDK4 are frequently upregulated, excessively active and overexpressed in cancer cells including breast, lung, and prostate cancers, and adrenocortical carcinomas(Baker et al. 2022; Lapenna and Giordano 2009; Liang et al. 2020; Malumbres and Barbacid 2009; Wu et al. 2011; Zhang et al. 2021a), making them promising drug discovery targets.(Baker et al. 2022; Lapenna and Giordano 2009; Liang et al. 2020; Wu et al. 2011; Zhang et al. 2021a) Inhibiting them can impede cell cycle progression, curtailing cell division and proliferation.(Fassl et al. 2022) The activity of CDK4 is often increased in cancer cells due to genetic mutations, amplifications, and overexpression of upstream nodes in the signaling pathways and in its regulatory cyclin-D. CDK4 plays a crucial role in the G1 phase–the longest phase of the cell cycle–making it an important target for cancer therapy.(Adon et al. 2021; DeMichele et al. 2015; Lim et al. 2016; Vijayaraghavan et al. 2018; Wander et al. 2022) CDK2, activated by cyclin-E binding in the G1/S transition also acts in DNA damage repair.(Tadesse et al. 2019) Significantly, although mutations in *CDK2* and *CDK4* genes have been identified in some cancer cells, different than other kinases such as PI3K(Galdadas et al. 2020), their frequency is low, especially in the catalytic domain, as shown in The Cancer Genome Atlas (TCGA) and Genomics Evidence Neoplasia Information Exchange (GENIE) databases,(Otto and Sicinski 2017) emphasizing the severe outcome of their mutations.

Numerous studies have shed light on the molecular mechanisms underlying the regulatory roles of CDKs in cell cycle progression and their inhibition by small molecules.(Arter et al. 2022; Fassl et al. 2022; Guiley et al. 2019; Herrera-Abreu et al. 2016; Hope et al. 2023; Tadesse et al. 2019; Whittaker et al. 2017; Zhang et al. 2021a) The CDK interacting protein/Kinase inhibitory protein (CIP/KIP) inhibitors bind to CDK2 complexes, hampering the kinase activity, while INhibitors of CDK4 (INK4) bind to CDK4 or CDK6, preventing their interaction with cyclin-D.(Fassl et al. 2022; Guiley et al. 2019; McGrath et al. 2017) CDK4 and CDK6 inhibitors have shown efficacy in treating estrogen receptor-positive breast cancer and are now approved for clinical use.(Herrera-Abreu et al. 2016)

In the canonical cell cycle activation pathway, CDK4 is activated in the G1 phase, while CDK2 is involved in the G1/S transition and throughout the S-phase.(Hume et al. 2020) Active CDK4 in complex with cyclin-D partially phosphorylates the retinoblastoma protein (Rb), disrupting the Rb/E2F interaction and releasing E2F transcription factors, which then trigger the expression of cyclin-E. Subsequent CDK2 activation by cyclin-E leads to hyperphosphorylation of the Rb protein.(Kim et al. 2022; Rubin et al. 2020)

While the sequences of CDK2 and CDK4 differ (**Fig. 1a**), they share the typical protein kinase conformation (**Fig. 1b**). The crystal structure of active CDK2 in complex with cyclin-E is available, while the active CDK4 structure in complex with cyclin-D has long been elusive. CDK4 retains a stable inactive conformation even when cyclin-bound and phosphorylated on the activation segment, suggesting that binding of the protein substrate and other factors are required for full activation (**Fig. 1c**).(Day et al. 2009) Gharbi et al.(Gharbi et al. 2022) have recently determined the crystal structure of active CDK4 in complex with cyclin-D. Recent studies using molecular dynamics (MD) simulations and other computational approaches reported differences in the conformational landscapes of CDK2 and CDK4 complexes as well as the effects of CDK inhibitors on the activation of these complexes.(Arora et al. 2023; Floquet et al. 2015; Swadling et al. 2022; Wood et al. 2019) These studies suggested that CDK2 and CDK4 have distinct conformational dynamics and allosteric interactions with their regulatory cyclins and peptide inhibitors, which may contribute to their different activation kinetics.

**Figure 1.**
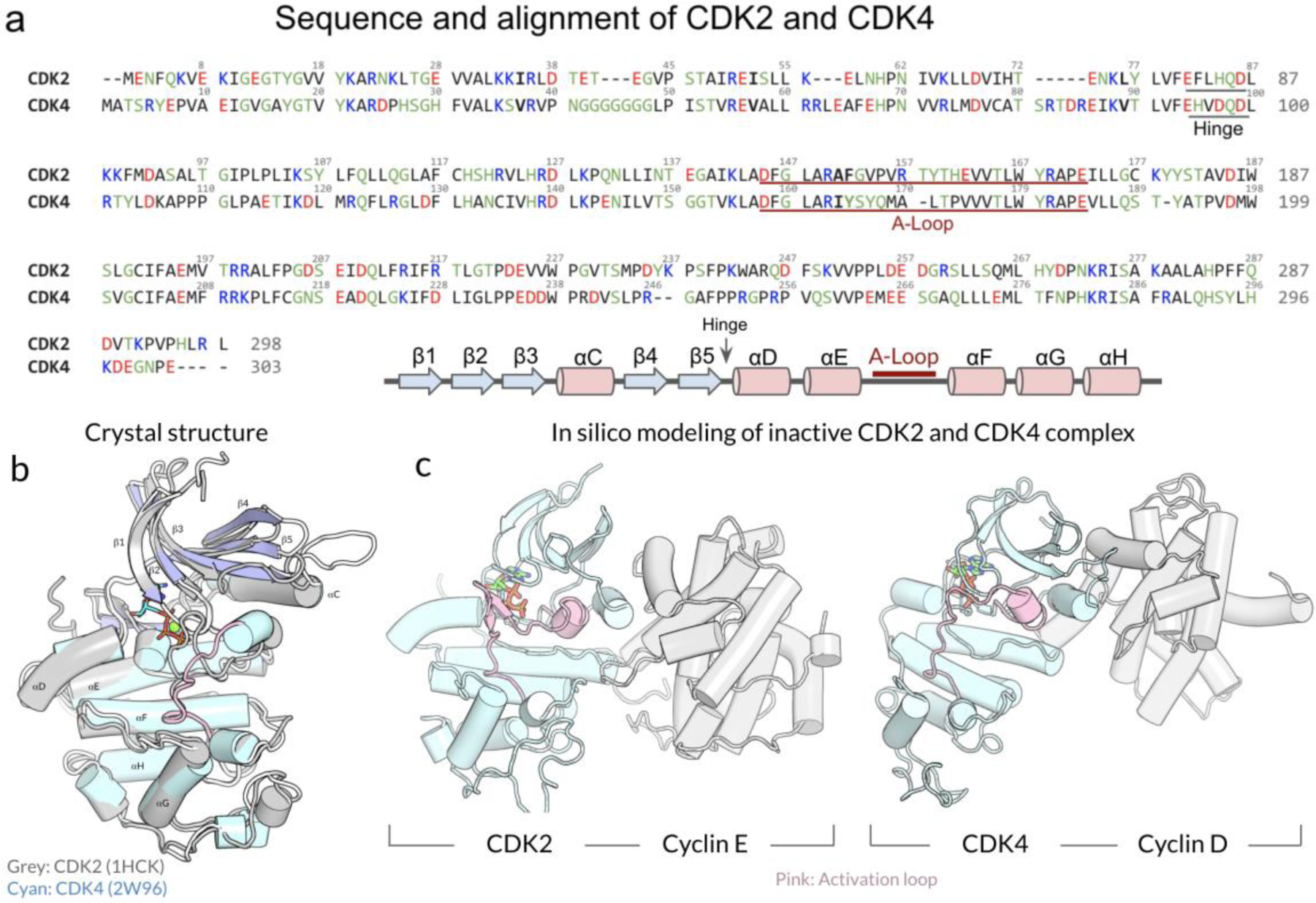
Sequence and structural alignments of CDK2 and CDK4. (a) Sequence and structural alignment of CDK2 and CDK4, showcasing their similarities and differences. (b) Superimposition of the crystal structures of CDK2 and CDK4. Key domains and residues are labeled. (c) *In silico* modeling of the inactive CDK2 and CDK4 in complex with their cognate cyclin partners, cyclin-E, and cyclin-D, respectively. The activation loops, crucial for kinase activity, are highlighted in pink.

Despite their vital importance and significant progress, the precise mechanisms by which CDK4 and CDK2 are activated by cyclin-D in the G1 phase and cyclin-E in the G1/S transition of the cell cycle, respectively, remain unresolved. The duration of the G1 and the G1/S transition in the cell cycle differ. G1 is long. G1/S is short. The expression levels and concentration of the respective cyclins differ as well. Cyclin-E expression level is regulated by a positive feedback loop, peaking, and falling over a short time span, which is not the case for cyclin-D. CDK4 has been traditionally deemed as the more accessible for drug discovery, hinting its potential for more detailed investigation. Thus, in this work, we aim to understand why cyclin-E/CDK2 have been recruited for the transition point and cyclin-D/CDK4 for the G1 phase. We query their conformational properties and landscapes and ask which states and activation pathways are favored. We focus on the inactive conformations of CDK2 and CDK4 in their monomeric state, as well as in complex with their respective cyclin partners (**Fig. S1**). Explicit solvent MD simulations of these systems were performed for the CDKs with and without ATP in the active site. Our idea has been to model the activation of cyclin-E/CDK2 and cyclin-D/CDK4 each step of way along the conformational energy landscape to determine the preferred sequence of the activation. This leads us to determine what we perceive as the fundamental guiding principle shaping the activation pathways of the two cell cycle systems: their rates of activation. Activation of cyclin-E/CDK2 which executes a cell cycle stage transition is fast, whereas that of cyclin-D/CDK4 is slow.

Here we analyze the conformational landscape of CDK2 and CDK4 upon ATP and cyclin binding and propose activation steps for these two CDK complexes. Our observations support CDK2 preferentially binding ATP first then cyclin-E, activating rapidly, while CDK4 favors binding cyclin-D first then ATP, resulting in a slower activation process. CDK2 has a more flexible activation loop (A-loop) and greater tendency to facilitate ATP loading compared to CDK4. CDK2 is also more responsive to ATP or cyclin binding compared to CDK4, suggesting a tendency to shift toward its active conformation more readily. We attribute their differences to longer CDK4 loops connecting the β3-strand to the αC-helix and the β4-strand to the β5-strand, which block the αC-helix from moving inward. A key driving force of CDK4 slower activation is the stronger hydrophobic interactions between the A-loop and the αC-helix. This is bolstered by the interaction of the longer CDK4 loops which, upon cyclin binding, thwart the formation of the catalysis-ready state compared to CDK2. Our studies contribute to a deeper understanding of the activation mechanisms, potentially informing the development of novel therapeutics.

## 2 | RESULTS

### 2.1 | CDK2 is more pre-organized compared to CDK4 to facilitate ATP loading

Protein kinases play crucial roles in regulating cellular processes by transferring phosphate groups from ATP molecules to specific protein substrates. The ATP-loading ability of the kinases is one of the determinants for their efficiency in catalysis. Kinases allowing easier ATP entry and more stable binding are more likely to efficiently catalyze reactions. The A-loop can either block or allow ATP entry and plays an important role in regulating the transfer of phosphate groups. In our simulations, the monomeric apo states of CDK2 and CDK4 (hereafter CDK2^Apo^ and CDK4^Apo^, respectively) adopted an inactive conformation with the collapsed A-loops, which obstructs access to both the ATP-binding pocket and substrate binding site.(Modi and Dunbrack 2019) For CDKs to effectively carry out their phosphorylation functions, the A-loops need to be repositioned, and the ATP-binding pocket must expand to accommodate ATP loading. Recent studies highlighted the flexibility of the A-loop and the dimensions of the ATP-binding pocket as crucial factors influencing the ability of CDKs to load ATP.(Chakraborty et al. 2019; Pucheta-Martinez et al. 2016) Building upon these findings, we assess the flexibility of the A-loops in both CDK2^Apo^ and CDK4^Apo^. Despite their structural similarities, they have differences in their A-loop sequences and dynamics (**Fig. 2a**). Superimposed A-loop conformations over the simulation trajectories illustrate that the monomeric CDK2^Apo^ has more flexible A-loop than the monomeric CDK4^Apo^. The root-mean-squared-fluctuations (RMSFs) verify that CDK2^Apo^ shows increased fluctuation of the A-loop as compared to CDK4^Apo^ (**Fig. 2b).** The different A-loop dynamics may be caused by its different length, which is longer in CDK2 than in CDK4. Based on our observations, CDK2 is more likely to extend its A-loop, allowing ATP entrance, which can explain why the recent NMR observations for the sampled CDK2^Apo^ conformations with the A-loop OUT state.(Majumdar et al. 2021) Our results show that CDK2^Apo^ is more pre-organized to facilitate ATP loading, suggesting a greater readiness for activation and substrate phosphorylation compared to CDK4.

**Figure 2.**
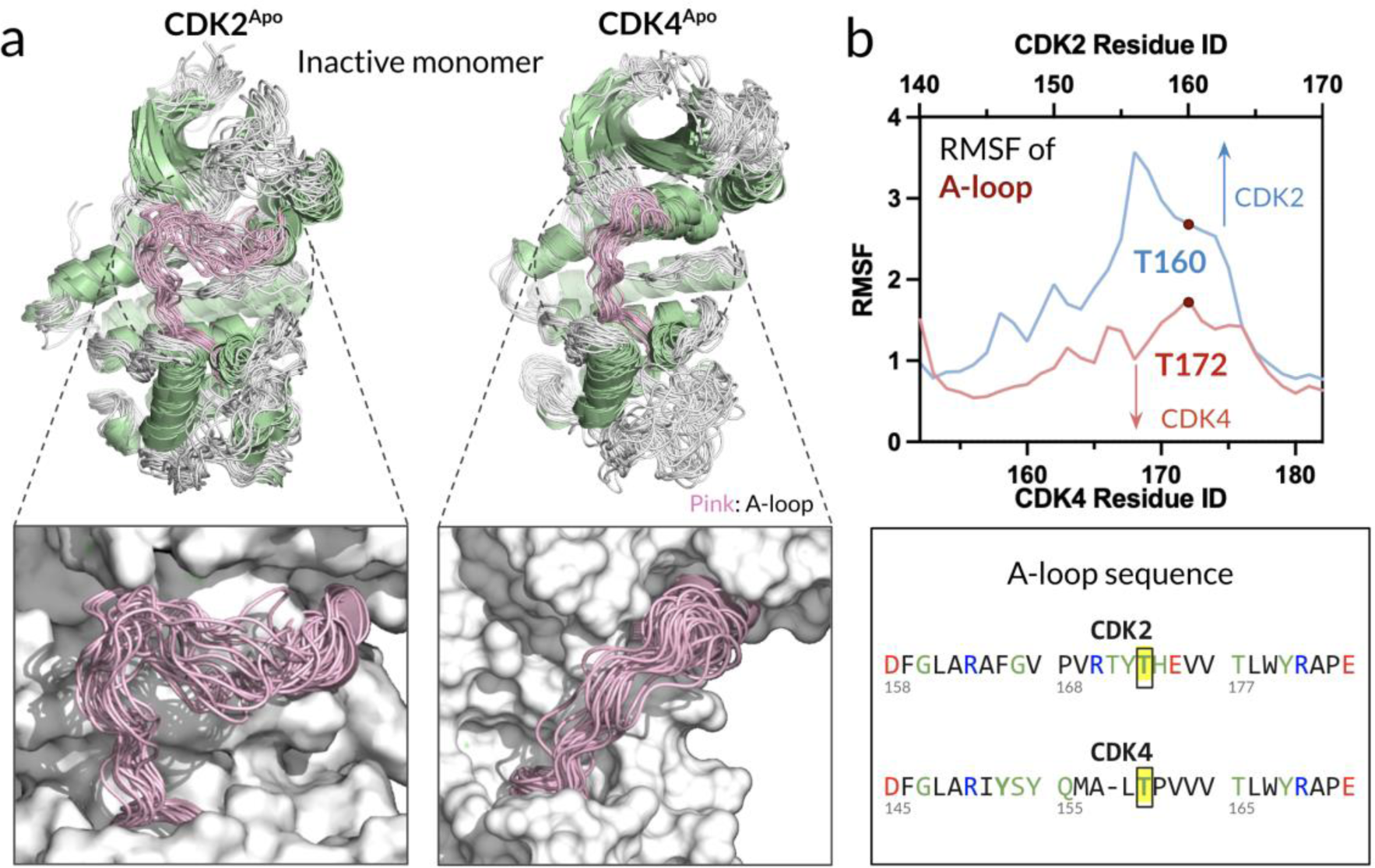
Activation loop of CDK2^Apo^ has a greater flexibility than CDK4^Apo^, facilitating ATP entrance. **(a)** Superimposition of snapshots of the monomeric inactive CDK2^Apo^ reveals that the A-loop of CDK2 exhibits greater flexibility as compared to CDK4. This increased flexibility may contribute to differences in ATP binding and activation kinetics between these two kinases. **(b)** Root-mean-squared-fluctuation (RMSF) and sequence alignment of the A-loops for CDK2 and CDK4. Yellow highlighted regions indicate the phosphorylation sites, which play a crucial role in modulating kinase activity. The RMSF values demonstrate the extent of flexibility within the A-loops, further illustrating the inherent structural differences between CDK2 and CDK4.

To compare the conformations of the ATP-binding pockets, the residue-based distance between the valine (V18 in CDK2 and V20 in CDK4) in the roof and the leucine (L134 in CDK2 and L147 in CDK4) in the bottom of the active-site cleft were calculated as a proxy for the ATP-binding pocket’s open/closed conformations. We observed that the monomeric CDK2^Apo^ has an open ATP-binding pocket as indicated by the peak at ∼15.4 Å in the V18-L134 distance distribution curve (**Fig. 3a**, blue curve), facilitating ATP entrance to the pocket. For the monomeric CDK2^ATP^, once ATP enters the active-site cleft, the pocket is closed. For the cyclin-E/CDK2^Apo^ complex, cyclin binding also yields the closed ATP-binding pocket, but less effectively than in the cyclin-E/CDK2^ATP^ complex. In contrast, the monomeric CDK4^Apo^ exhibits a closed ATP-binding pocket as indicated by the peak at ∼13.5 Å in the V20-L147 distance distribution curve (**Fig. 3b**, blue curve), making spontaneous ATP loading less likely. To load ATP, CDK4 requires cyclin-D binding that shifts the equilibrium toward the open ATP-binding pocket. For the cyclin-D/CDK4^Apo^ complex, a shift of the peak in the distance distribution curve from ∼13.5 Å to ∼14.5 Å is observed. Once ATP enters the active-site cleft, the cyclin-D/CDK4^ATP^ pocket closes as evident from a growing peak at ∼13 Å in the V20-L147 distance distribution curve. Our findings also indicate that the monomeric CDK2^ATP^ has a higher affinity for the ATP compared to CDK4^ATP^ (**Fig. S2)**. In complex with cyclin, CDK2 has a slightly higher affinity for ATP than CDK4, suggesting that CDK2 prefers an ATP-bound state, while CDK4 prefers an ATP-free state. This observation aligns with the fact that a crystal structure of monomeric CDK2 with ATP is available, while no such crystal structure is currently available for CDK4. This may support higher CDK2 substrate phosphorylation efficiency than CDK4, indicating that CDK2 more readily shifts its conformation toward its active state and stabilizes the ATP.

**Figure 3.**
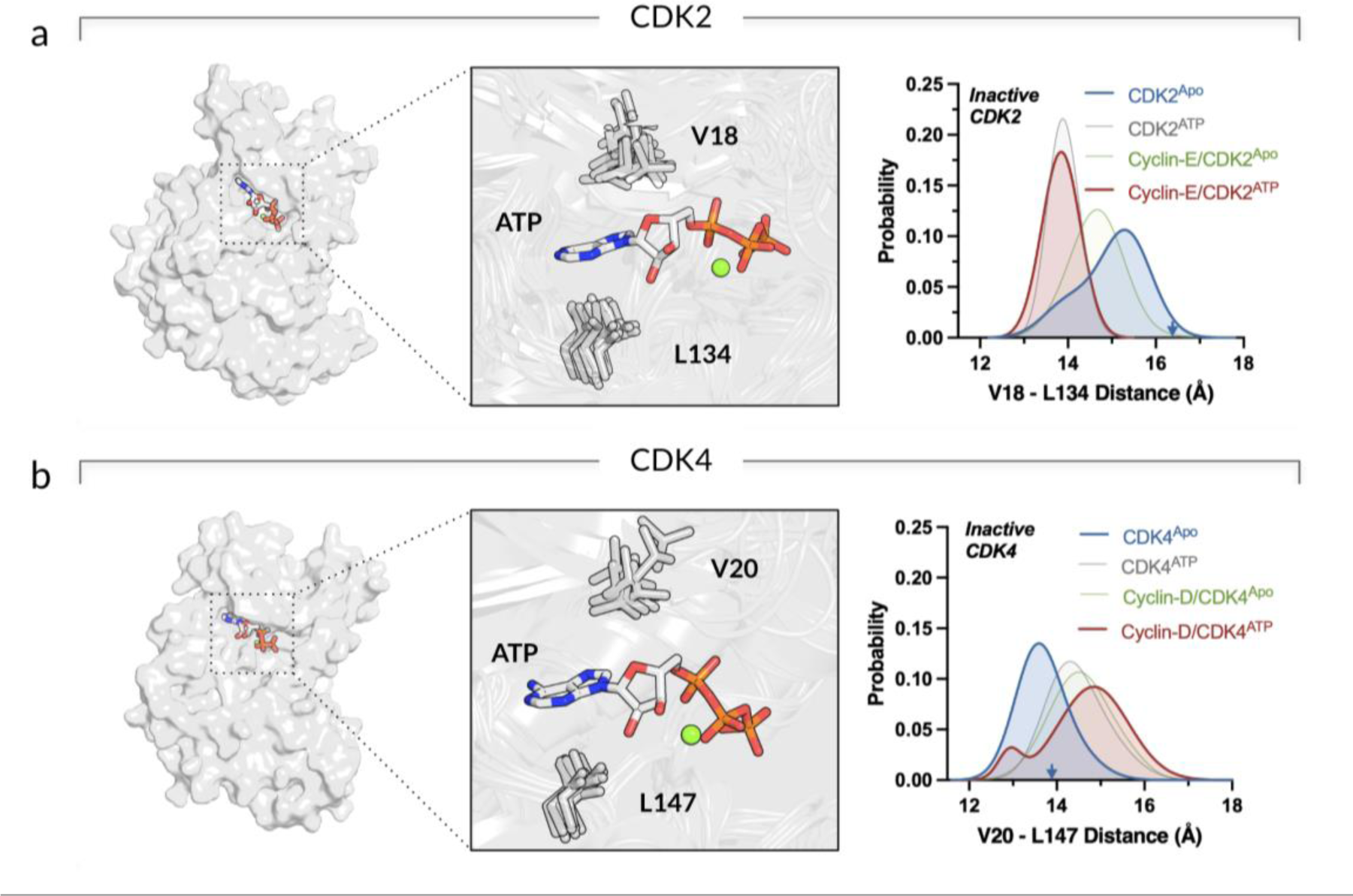
Opening/closing ATP-binding pocket for CDK2 and CDK4 upon ATP loading and cyclin binding. **(a)** Surface representation of CDK2 with ATP in the active site (*left panel*). Highlight showing the active site (*middle panel*). Probability distribution functions of the distance between V18 in the roof of the active-site cleft and L134 in the bottom of the active-site cleft for CDK2^Apo^ (blue), CDK2^ATP^ (gray), cyclin-E/CDK2^Apo^ (green), and cyclin-E/CDK2^ATP^ (red) (*right panel*). **(b)** The same for the CDK4 systems with the active site residues, V20 and L147 (*left and middle panels*). The probability distribution functions show the V20-L147 distance for CDK4^Apo^ (blue), CDK4^ATP^ (gray), cyclin-D/CDK4^Apo^ (green), and cyclin-D/CDK4^ATP^ (red) (*right panel*).

### 2.2 | ATP or cyclin binding facilitates a more accessible shift to the active conformational ensemble in CDK2 compared to CDK4

Protein kinase activation involves conformational changes of the αC-helix and the A-loop. Inspection of the dynamics of the two motifs is essential for monitoring the kinase conformational transitions. As a typical protein kinase, CDKs adopt an IN αC-helix in the active state and an OUT αC-helix in the inactive state.(Morgan 2007) The transition from the inactive to the active conformation involves ∼30° rotation of the αC-helix.(De Bondt et al. 1993) This rotation results in forming a conserved salt bridge between a glutamic acid (E51 in CDK2 and E56 in CDK4) in the αC-helix and a lysine (K33 in CDK2 and K35 in CDK4) in the β3-strand. The salt bridge stabilizes the IN αC-helix and the active conformation. In the catalytic active state, the salt bridge stabilizes the γ-phosphate of ATP for phosphate transfer, while in the inactive state, the salt bridge is disrupted, inhibiting the catalytic reaction.(Arter et al. 2022; Malumbres 2014) To monitor the conformational dynamics of the αC-helix in inactive CDKs, we calculated the residue pair distance for the salt bridge. We observed that the αC-helix of inactive CDK2 has a greater tendency to move inward upon ATP loading and cyclin binding (**Fig. 4a**). The monomeric CDK2^Apo^ retained the K33-E51 distance at ∼17.9 Å as in the crystal structure. However, for the monomeric CDK2^ATP^, the K33-E51 distance decreased upon ATP loading (blue curve shifting to grey curve), and for cyclin-E/CDK2^Apo^ upon cyclin-E binding (blue curve shifting to green curve). For cyclin-E/CDK2^ATP^ with both ATP loading and cyclin-E binding, the distance further decreased to ∼15 Å (red curve), suggesting that either ATP loading or cyclin-E binding drives CDK2 towards the active state. In marked contrast, the conformational dynamics of the αC-helix of inactive CDK4 is less responsive to both ATP loading and cyclin-D binding (**Fig. 4b**). The monomeric CDK4^Apo^ yielded the K35-E56 distance at ∼14.8 Å. ATP loading (CDK4^ATP^) and cyclin-D binding (cyclin-D/CDK4^Apo^), induce a smaller change in the K35-E56 distance, suggesting that additional factors may be needed to facilitate activation. The K35-E56 distance of ∼14.8 Å in the CDK4^Apo^ is not as far apart as the corresponding K33-E51 distance of ∼17.9 Å in the CDK2^Apo^, indicating that the αC-helix of CDK4 appears to have limited outward movement as much as that of CDK2. We attribute these observations to the structural and sequence differences between CDK2 and CDK4. In CDK4, the loops connecting the β3-strand to the αC-helix (loop^β3-aC^) and the β4-strand to the β5-strand (loop^β4-β5^) are longer than those in CDK2 (**Fig. 4c**). Interestingly, we observed that the longer loops of CDK4, loop^β3-aC^ and loop^β4-β5^ interact with each other, preventing the αC-helix from moving inward.

**Figure 4.**
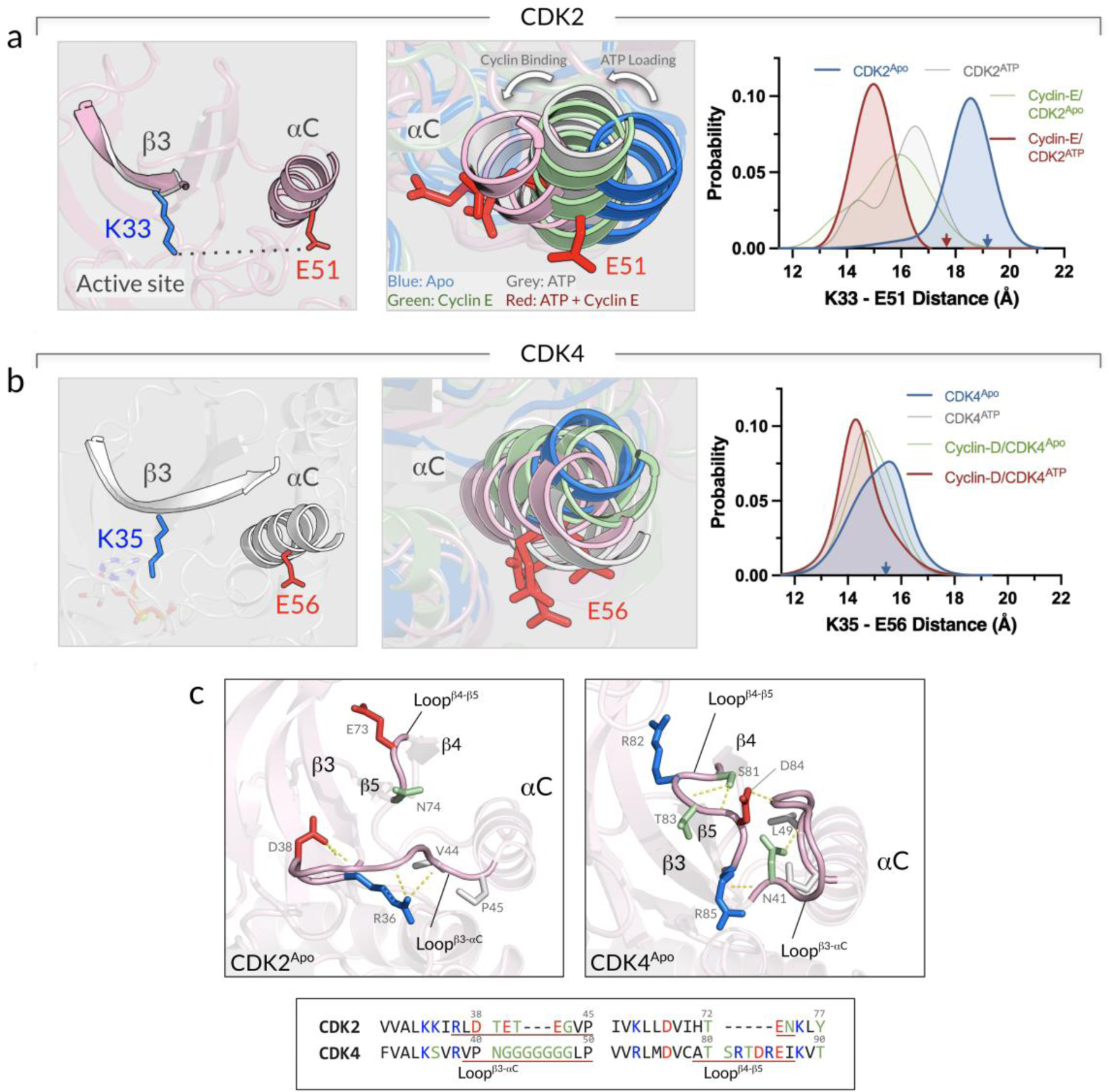
ATP loading and cyclin binding induce changes to the αC-helix position of CDK2 compared to CDK4. **(a)** Snapshot representing the salt bridge residues pair, K33 on the β3-strand and E51 on the αC-helix of CDK2 (*left panel*). Superimposition of the αC-helices of the CDK2 systems (*middle panel*). Probability distribution functions of the K33-E51 distance for CDK2^Apo^ (blue), CDK2^ATP^ (gray), cyclin-E/CDK2^Apo^ (green), and cyclin-E/CDK2^ATP^ (red) (*right panel*). **(b)** The same for the CDK4 systems with the salt bridge residues pair, K35 and E56 (*left and middle panels*). The probability distribution functions show the K35-E56 distance for CDK4^Apo^ (blue), CDK4^ATP^ (gray), cyclin-D/CDK4^Apo^ (green), and cyclin-D/CDK4^ATP^ (red) (*right panel*). **(c)** Snapshots showing the conformations of the loops connecting the β3-strand to the αC-helix and the β4-strand to the β5-strand for CDK2^Apo^ (*left panel*) and CDK4^Apo^ (*right panel*). These two loops in CDK4 are longer than those in CDK2 (*bottom panel*), showing extensive interactions and blocking the αC-helix from moving inward.

The coupling of the A-loop and the αC-helix in CDK2 and CDK4 shows a contrasting behavior. To illustrate this, we examined the conformational changes of the A-loops to determine how the ATP loading or cyclin binding affects the kinase conformation toward the active state. The representative snapshots of the CDK systems highlight the conformational changes of the A-loop and its coupling with the αC-helix (**Fig. S3**). To evaluate the different A-loop conformations, we calculated the distance between the αC-helix and the αL12-helix of each CDK. The αL12-helix is a small helix in the A-loop and is spatially adjacent to the αC-helix of the CDKs in the inactive state (**Fig. S4**). For the CDK2 systems, the αL12-helix moves away from the αC-helix as the αC-helix moves inwards upon ATP loading (CDK2^ATP^) or cyclin-E binding (cyclin-E/CDK2^Apo^) (**Fig. 5a)**. The αL12-helix moves farther away from the αC-helix in the cyclin-E/CDK2^ATP^ complex, suggesting that the A-loop has a tendency to extend upon both ATP loading and cyclin-E binding. The outward movement of the αL12-helix destabilizes this inhibitory helix, leading to an extended A-loop as in the active form of CDK2 (**Fig. S5**). We suspect that the movements of the A-loop and αC-helix are coupled in CDK2 on its way to activation. For the CDK4 systems, however, the αL12-helix slightly moves toward the αC-helix as the αC-helix moves inwards upon ATP loading (CDK4^ATP^) and cyclin-D binding (cyclin-D/CDK4^Apo^) (**Fig. 5b**). The αL12-helix is closer to the αC-helix in the cyclin-D/CDK4^ATP^ complex. This contrasting CDK4 behavior can be explained by the αL12-helix of CDK2 having weaker hydrophobic interactions with the αC-helix, with A151 in the αL12-helix compared to the I164 of CDK4. Recent experiments using double electron-electron resonance (DEER) spectroscopy revealed that wild-type CDK2 can populate the ensembles of the A-loop OUT conformation, but CDK2 with the A151I mutation (CDK4-specific) causes the ensembles to shift in the absence of the A-loop OUT conformation.(Majumdar et al. 2021) CDK2 with ATP and cyclin-E favors the A-loop OUT conformation, which eliminates the inhibitory helix and facilitates full A-loop extension. In contrast, CDK4 with ATP and cyclin-D retains the inhibitory helix that interacts with the αC-helix, stabilizing the inactive state. Thus, we hypothesize that CDK2 has a more dynamic and flexible A-loop movement than CDK4, which confers a kinetic advantage for its activation.

**Figure 5.**
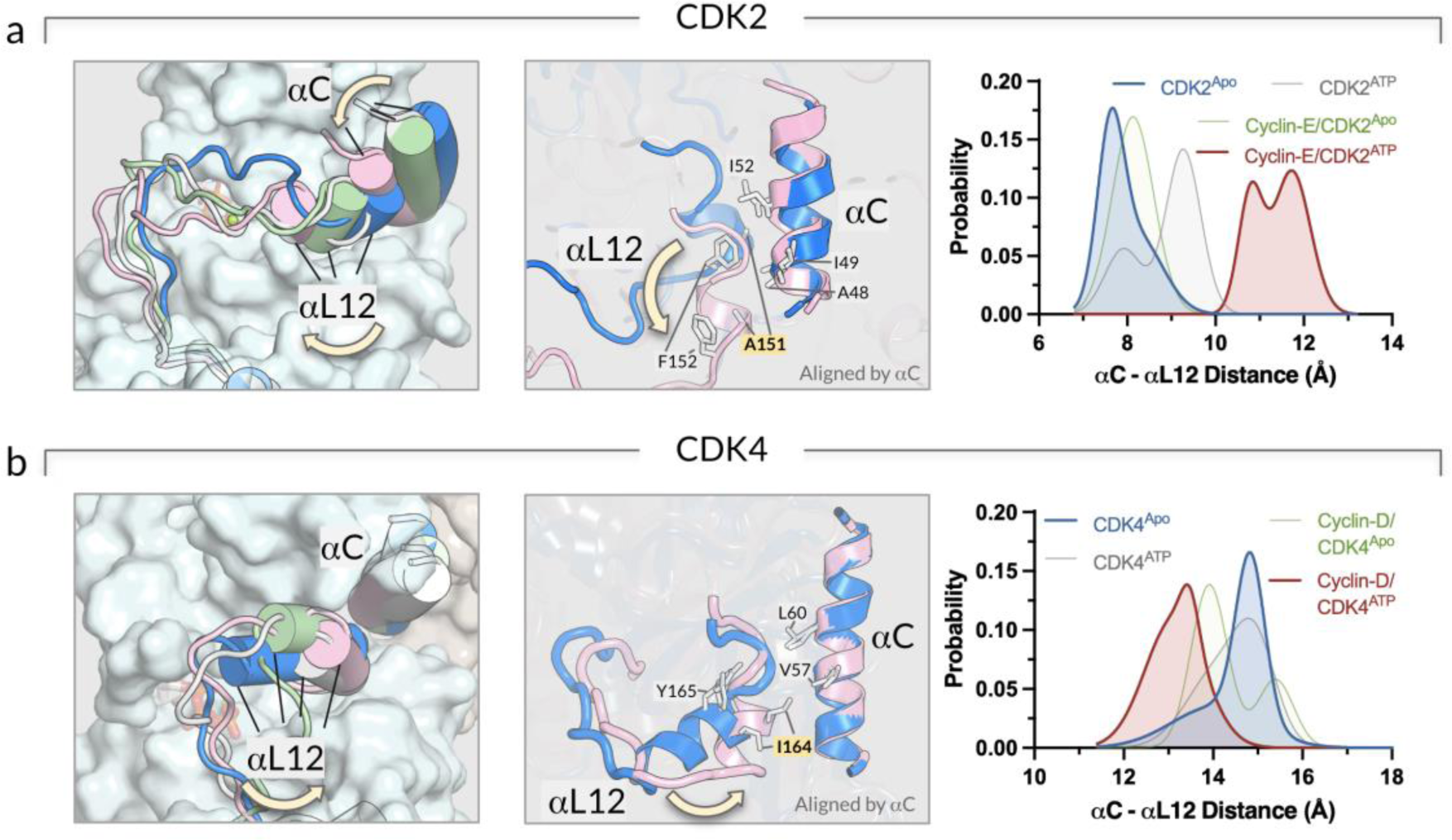
The coupling between the A-loop conformation and the αC-helix is stronger in CDK2 than CDK4. **(a)** Snapshots representing the superimposed conformations of the A-loop of the CDK2 systems (*left panel*). Structural alignment of the A-loop of the ATP-loaded CDK2 in complex with cyclin-E (red cartoon) with respect to apo CDK2 (blue cartoon) (*middle panel*). Probability distribution functions of the αC-αL12 distance for CDK2^Apo^ (blue), CDK2^ATP^ (gray), cyclin-E/CDK2^Apo^ (green), and cyclin-E/CDK2^ATP^ (red) (*right panel*). **(b)** The same for the CDK4 systems (*left and middle panels*). The probability distribution functions show the αC-αL12 distance for CDK4^Apo^ (blue), CDK4^ATP^ (gray), cyclin-D/CDK4^Apo^ (green), and cyclin-D/CDK4^ATP^ (red) (*right panel*). Yellow arrows highlight the direction of the movement of αL12-helix.

### 2.3 | Differential activation mechanisms of cyclin-E/CDK2 and cyclin-D/CDK4 via different binding preferences for ATP and cyclin partners

We decipher the activation mechanisms of cyclin-E/CDK2 and cyclin-D/CDK4, elucidating the preferred order in which these kinases bind to their respective cyclin partners and ATP molecules.

Above, we presented the conformational changes of the ATP-binding pocket upon cyclin binding to see how cyclin stabilizes the pocket for efficient phosphorylation. For CDK2, ATP loading onto CDK2^Apo^ is highly favored due to the open pocket. However, cyclin-E binding to CDK2^Apo^ causes the pocket to close, reducing the probability of ATP loading. Thus, the most populated pathway for CDK2 activation involves ATP loading as the initial step. Binding of cyclin-D to CDK4^Apo^ results in the opening of the ATP-binding site, promoting subsequent ATP loading. However, ATP loading on CDK4^Apo^ is less likely due to the closed ATP-binding pocket. Thus, the most populated pathway for CDK4 is to first bind cyclin-D. Taken together, we suggest that inactive CDK2 preferentially loads ATP first and then binds cyclin-E (**Fig. 6**). Conversely, inactive CDK4 prefers to bind to cyclin-D first, which opens the ATP-binding pocket and facilitates ATP loading. In both cases, the ATP-bound inactive cyclin/CDK complexes undergo significant conformational changes, such as the αC-helix adopting an IN position and the A-loop extending outward to interact with their cyclin partners.

**Figure 6.**
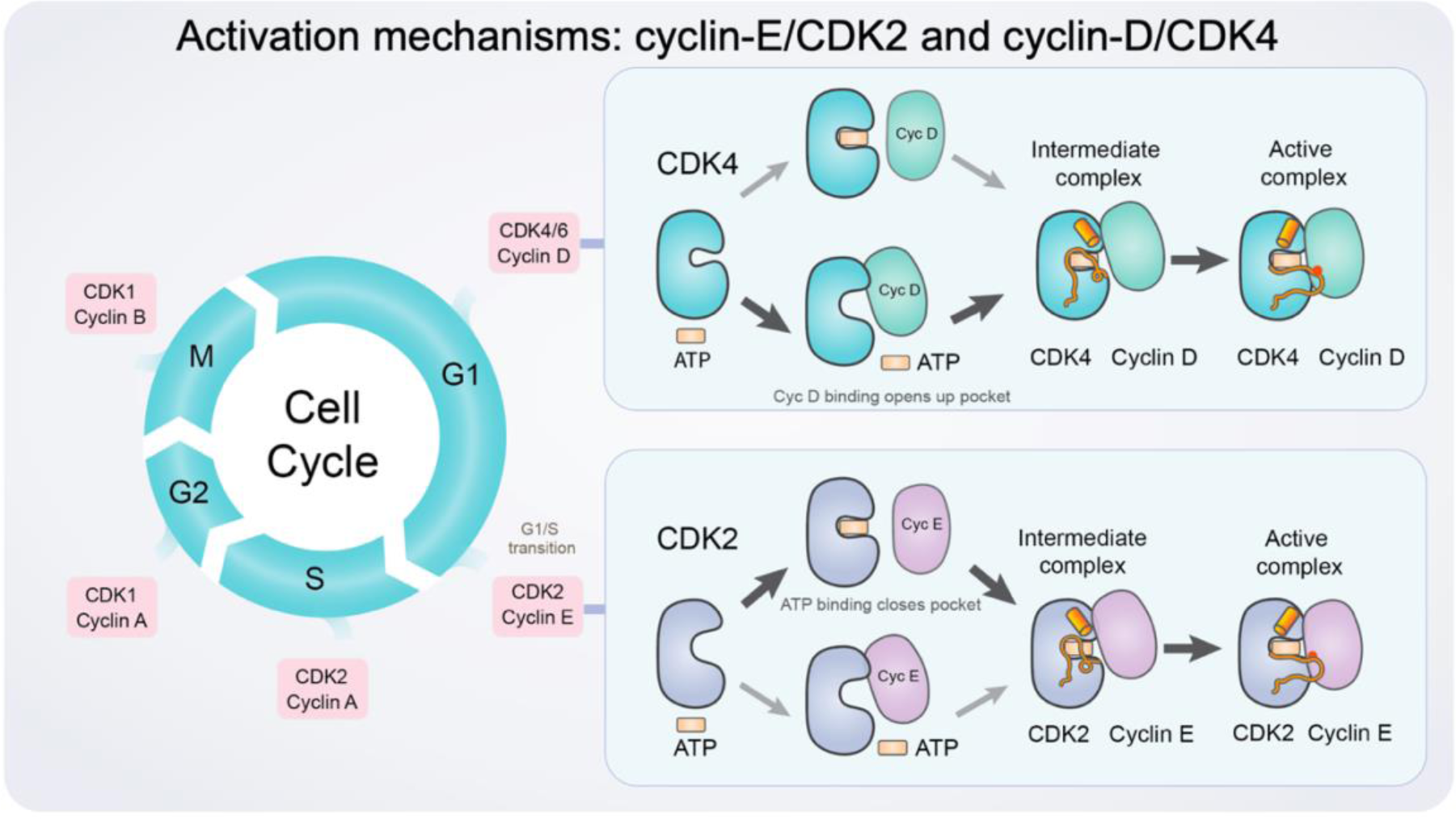
A schematic diagram illustrating the activation mechanism of cyclin-E/CDK2 and cyclin-D/CDK4. Cyclins and CDKs in the cell cycle (*left panel*). Cell cycle progression is regulated by complexes of CDKs and their cyclin partners. The key regulatory steps and protein interactions contributing to the activations of CDK2 and CDK4 (*right panel*). The ATP-binding pocket of CDK2^Apo^ is open and flexible, facilitating ATP loading. Upon ATP loading, the pocket is closed, and cyclin-E binding promotes the active kinase conformation. In contrast, the ATP-binding pocket of CDK4^Apo^ is closed and rigid, impeding ATP entrance to the pocket. Cyclin-D binding induces the open conformation of the pocket, allowing ATP to load. Orange tubes are A-loops; cylinders are αC-helix; red dots are the phosphorylation sites on the A-loop. The major activation pathway for each CDK is indicated by bold arrows.

## 3 | DISCUSSION

Cyclin-D/CDK4 and cyclin-E/CDK2 play distinct roles in the cell cycle. Here, we ask *why* cyclin-D/CDK4 preferentially acts in the G1 phase and cyclin-E/CDK2 in the G1/S transition. We also ask *how* they were optimized for their distinct functions. Our observations lead us to postulate that cyclin-D/CDK4 preferentially executes the G1 phase and cyclin-E/CDK2 the G1/S transition because cyclin-E/CDK2 has a faster activation time than cyclin-D/CDK4, and we detail how. Our innovative concept is based on observations from our data, cell biology, and experimental structural work.

### 3.1 | The distinct mechanisms of activation of cyclin-D/CDK4 and cyclin-E/CDK2

In our scenarios, CDK2 is pre-organized to allow ATP loading prior to cyclin-E binding. In contrast, CDK4 exhibits a preference for cyclin-D binding before ATP loading. In line with this, (i) upon ATP loading and cyclin binding, the αC-helix of CDK2 experiences a significant inward movement, toward the active state. In contrast, the αC-helix of CDK4 remains relatively static in the inactive form, regardless of ATP loading or cyclin binding, suggesting a slower conformational change. (ii) The A-loop and the αC-helix of CDK2 are coupled, meaning that the inward movement of αC is correlated with A-loop extension. However, in CDK4, the A-loop and the αC-helix are more rigid and resist movement, requiring more energy to switch between states. (iii) CDK2 has a more flexible and responsive ATP-binding site than CDK4, providing a kinetic advantage for its activation. Significantly, (iv) in the cell, CDK2 activation is part of a positive feedback loop, resulting in switch-like behavior, while CDK4 activation is a more gradual process. In the G1 phase, active CDK4 (and CDK6) in complex with cyclin-D phosphorylates the Rb protein, triggering a partial release of the E2F transcription factor, which in turn induces cyclin-E expression. Cyclin-E activates CDK2, which further phosphorylates Rb, leading to the complete release of E2F, thereby enhancing cyclin-E expression further activating CDK2, and promoting hyperphosphorylation of Rb.(Morgan 2007; Rubin et al. 2020) This canonical pathway demonstrates that CDK2 activation during the G1 phase to G1/S transition is part of a positive feedback loop. Recent studies proposed that cyclin-D/CDK4 complex can either directly hyperphosphorylate Rb, leading to cell cycle progression into the S phase(Chung et al. 2019; Yang et al. 2020), or indirectly activate CDK2 by sequestering KIP/CIP family protein inhibitors away from CDK2, allowing CDK2 to hyperphosphorylate Rb and promote the cell’s entry into the S phase.(Narasimha et al. 2014) Further, significantly, (v) in terms of duration, CDK4 primarily governs progression through the G1 phase, which is the longest phase of the cell cycle, lasting approximately 11 hours in rapidly proliferating human cells.(Cooper 2000) In contrast, CDK2 regulates the G1/S phase transition, a relatively short period during which the cell decides whether to commit to DNA replication or not, as there is no turning back. Consequently, the activation time of CDK2 is shorter than that of CDK4, further highlighting the dynamic and time-sensitive nature of this critical phase in the cell cycle. In addition, (vi) related to the dynamics of the cell cycle, are the concentrations of cyclins in the multiple phases of the cell cycle, which have been assembled from multiple sources.(Nussinov et al. 2021)

### 3.2 | Experimental support for the proposed mechanisms

Experimental structural reports point to (vii) differences in the crystal structures of cyclin-E/CDK2 and cyclin-D/CDK4. They indicate that cyclin-E/CDK2 has a more stable active conformation. Proteins can adopt various conformations, and their crystal or cryo-EM structures only capture snapshots of these.(Bai et al. 2015; Boehr et al. 2009; Stehle et al. 2023; Xie et al. 2020) The most likely scenario is that the conformation captured in the crystal structure represents the one with the lowest energy under the crystallization conditions. A comparison of the available crystal structures of CDK2 and CDK4 provides evidence for differences in their activation dynamics. CDK2 has been observed in the active state (αC-in and A-loop extended) when complexed with cyclin-E.(Day et al. 2009; Takaki et al. 2009) In contrast, a crystal structure of fully active cyclin-D/CDK4 complex was not available until late 2022 when it was published by Gharbi et al.(Gharbi et al. 2022) This observation suggests that CDK2 has a more stable active conformation, implying faster and more efficient activation. The higher stability of the active CDK2 conformation in complex with cyclin-E, as opposed to CDK4, may reflect a lower energy barrier for the conformational changes required for activation. This lower energy barrier and faster activation kinetics would allow CDK2 to transition more rapidly from an inactive to an active state during the critical "point of no return" for committing to the DNA replication S phase. This difference in activation dynamics could have functional implications for the regulation of cell cycle progression and response to extracellular signals. Thus, the comparison of the crystal structures when considering their associated conformational potential energy landscapes supports the argument that cyclin-E/CDK2 is faster to activate than cyclin-D/CDK4. Lastly, the structural differences in the phosphorylated cyclin-E/CDK2 and cyclin-D/CDK4 complexes on the A-loop also suggest that the CDK2 complex activates faster than the CDK4 complex. Upon binding cyclin-E, CDK2 adopts an active conformation, in which the active site on the A-loop is buried in the complex.(Schachter et al. 2013; Takaki et al. 2009) This intrinsic conformational stability also hinders phosphatases from accessing the active site and dephosphorylating CDK2. In contrast, when bound to cyclin-D, CDK4 exhibits an exposed active site on the A-loop,(Day et al. 2009; Takaki et al. 2009) which facilitates fine-tuning of CDK4 activation by growth factors signaling.(Schachter et al. 2013) Although this dynamic regulation provides a higher degree of adaptability, it may result in slower activation due to the active state being continually modulated by phosphorylation and dephosphorylation.(Merrick et al. 2008)

### 3.3 | Our proposed mechanisms can explain why cyclin-E/CDK2 works at the sharp transition between G1 and S while cyclin-D/CDK4 is active during the G1 phase

Here we perform 2 μs simulations of the CDK2 and CDK4 (starting in their OFF states) with and without the appropriate cyclins in the presence and absence of ATP to investigate the intrinsic conformational propensities of each kinase. During the simulations, we observe that (i) CDK2 is structurally more poised to bind ATP due to flexible A-loop and (ii) more open nucleotide binding cleft as compared to CDK4. We also see that (iii) CDK2 more readily switches into the ON state (as indicated by the tendency of αC-helix position) upon nucleotide and/or cyclin binding as compared to CDK4. Lastly, (iv) we observe a more dynamic αL12-helix in the CDK2 which may allow for easier switching to the ON state of CDK2 as compared to CDK4. Taken together, CDK2 more readily binds ATP and is more readily switched into the ON conformation than CDK4. These observations lead us to suggest that these differences may help explain why CDK2 works at the sharp transition between G1 and S while CDK4 is active during the G1 phase. These observations need however to be put in the methodological and broader cellular context.

### 3.4 | Considerations of the timescales and the biology

Conformational transitions in kinases can happen at the millisecond and slower timescales, which are not captured in microsecond simulations. The timescale of the transitions observed here, and cell cycle transitions, are orders of magnitude different(Min et al. 2020), and the biological picture is fundamentally complex. Both kinases interact with CIP and Ink proteins, and recent work showed that the phosphorylation state of CIPs play an important role in CDK4 activation(Guiley et al. 2019). When interacting with the molecular chaperone Hsp90/Cdc37, CDK4 populates a re-arranged conformation(Verba et al. 2016), much beyond that observed in the simulations. Thus, the biological timing and function of CDK2 and CDK4 appear largely driven by deep energy wells created through interactions with cellular proteins and chaperones. Hsp90 can efficiently modulate the allosteric interactions and long-range communications for client protein activation.(Verkhivker 2022) At the same time, as we have shown here, the energetic fluctuations captured in the simulations are critical at the protein level, beyond that in the cellular environment.

Finally, the major contributor to fast CDK2 activation is the feedback mechanism between CDK2 and p21. A relatively small number of active CDK2 molecules can rapidly trigger p21 degradation releasing a burst of already active kinase molecules at the G1/S transition. (Cayrol and Ducommun 1998)

### 3.5 | Implications to drug discovery

Our studies offer insights into the conformational changes involved in cyclin-E/CDK2 and cyclin-D/CDK4 activation and their implications for drug discovery, particularly for CDK4, which is often targeted in cancer. CDK4 is overactive in cancer cells(An et al. 1999; Dai et al. 2016), and has a role in cell cycle (dys)regulation(Malumbres et al. 2004), making it an attractive drug target. Our findings of the distinct activation mechanisms of cyclin-E/CDK2 and cyclin-D/CDK4 suggest potential strategies for designing selective allosteric inhibitors for CDK4. The gradual activation process of CDK4 provides a longer time frame for inhibitors to bind, in contrast to CDK2. One possible approach for CDK4 allosteric inhibitors is stabilizing the inactive conformation, a classic allosteric kinase inhibitor design principle. Design strategies could also involve targeting conformational changes in CDK4 activation or blocking cyclin-D binding to CDK4. Allosteric inhibitors offer an alternative to ATP-competitive inhibitors, as they can be less toxic and more selective. Recent developments in allosteric inhibition of CDK2, targeting the interface between CDK2 and its cyclin partner(Betzi et al. 2011), provide promising examples for CDK4 inhibitor design.(Zhang et al. 2022) Accounting for the dynamic nature of protein binding sites and ATP pocket geometry is crucial for effective kinase inhibitor development.(Zhang et al. 2023) As to orthosteric drugs, as the recent crystal structure of CDK11 bound to the selective inhibitor OTS964 showed, evolutionary variations in the kinase domain can be exploited for drugs for closely related kinases.(Shrestha et al. 2022)

## 4 | CONCLUSIONS

The cell cycle plays a cardinal role in cell life and death. Under normal physiological conditions, it is carefully orchestrated; when dysregulated, it can lead to uncontrolled proliferation and cancer, to oncogene-induced senescence (OIS), as well as aberrant differentiation in neurodevelopmental disorders.(Nussinov et al. 2023) Cell cycle regulation entails complex processes. Full understanding requires grasp of its cellular and its structural biology. Several CDKs (e.g., CDK1, CDK2, CDK4 and 6) and cyclins (e.g., A, B, D, E) are involved in these processes. Despite their structural similarities and functional overlap, specific complexes are favored in distinct cell cycle stages: cyclin-D/CDK4/6 in the G1, cyclin-E/CDK2 in the G1/S transition; cyclin-A/CDK2 in the S stage; cyclin-A/CDK1 in the G2; and cyclin-B/CDK1 in the M stage. Our challenging aim is to figure out the structural basis of the cyclin/CDK preferences for the distinct cell cycle stages and merge these with cell data on protein function and regulation. We also aim to determine key structural characteristics that influence the catalytic efficiencies of these CDK complexes. To understand the workings of the cell these should be integrated with the structural changes of the protein complexes and the biology.

Within this framework, here we connected the mechanisms of activation of cyclin-D/CDK4 and cyclin-E/CDK2 to their functions in the G1 phase and G1/S transition in the cell cycle. These led us to postulate slower activation of cyclin-D/CDK4 and faster activation of cyclin-E/CDK2. We observed that cyclin-E/CDK2 prefers ATP loading prior to cyclin-E binding, while CDK4 prefers cyclin-D binding before ATP loading. We further observed that cyclin-E/CDK2 has a faster activation time than cyclin-D/CDK4 due to the conformational changes involved in its activation pathway, its flexible and responsive ATP-binding site, and the readily accessible stable active conformation. We also highlighted the importance of considering the conformational energy landscape for understanding *how* the activation mechanisms of cyclin-E/CDK2 and cyclin-D/CDK4 integrate with cell biology to accomplish their biological roles. Overall, these innovative mechanistic findings decipher overlooked hallmarks of the regulation of cell cycle progression, with potential implications for drug development. To our knowledge, our work pioneers the connection between the cyclins-CDKs mechanisms and their roles in cell cycle function.

Our study suggests that (i) the activation dynamics of CDK2 can be incorporated into a cellular expression level-based model of CDK2 activation, and that (ii) the mechanisms of CDK4 and CDK2 are distinct as well. Notably, the mechanism of activation of CDK1 and CDK2 are known to be distinct.(Merrick et al. 2008) Finally, (iii) our work raises the question of whether observations made here for CDK4 apply to CDK6, where different activation mechanisms have been proposed.(Bockstaele et al. 2009)

Our work provides an unprecedented mechanistic understanding of the distinct activation scenarios of cyclin-D/CDK4 and cyclin-E/CDK2 in cell cycle regulation, underpinning the slower activation of cyclin-D/CDK4 in the more extended G1 phase and the rapid activation of cyclin-E/CDK2 in the brief G1/S transition. Leveraging a range of experimental data and molecular dynamics simulations, we elucidate the inherent conformational dynamics and activation pathways of these key cell cycle regulators. Our findings not only address a long-standing question in cell cycle biology but also the design of more targeted and effective inhibitors against CDK4, opening new venues in cancer treatment.

## 5 | MATERIALS AND METHODS

### 5.1 | Modeling of inactive CDKs and CDKs complexes

We modeled the inactive conformations of CDK2 and CDK4 in their monomeric and cyclin-bound states in the presence and absence of ATP. The initial coordinates for the inactive CDK2 monomers were taken from the crystal structures of the ATP-bound state (PDB ID: 1HCK) and the apo state (PDB ID: 1HCL) (SchulzeGahmen et al. 1996), which generate monomeric inactive CDK2^ATP^ and CDK2^Apo^, respectively. The inactive cyclin-E/CDK2 complexes were modeled by replacing the inactive CDK2 with the active cyclin-E/CDK2 complex (PDB ID: 1W98).(Honda et al. 2005) For CDK4 systems, the initial coordinates for the inactive CDK4 monomers were extracted from the crystal structure of the inactive cyclin-D/CDK4 complex (PDB ID: 2W9Z)(Day et al. 2009). We simulated all systems in the presence and absence of ATP in the active site. The simulations are listed in **Table S1** and snapshots of all systems are in **Fig. S1**. We used the TIP3P water model and added Na^+^ and Cl^−^ to neutralize the solvated systems and maintain a physiological salt concentration of 150 mM.

### 5.2 | MD simulation protocol

We employed molecular dynamics (MD) simulations following the protocol established in our previous publications.(Jang et al. 2021; Jang et al. 2020; Liu et al. 2022a; Liu et al. 2022b; Zhang et al. 2021b) The simulations consisted of several stages. We began by conducting 10,000-step energy minimizations using the conjugate gradient minimization method to construct the systems. This step was crucial for eliminating unfavorable contacts between atoms in the systems. For each system, we performed a 2 μs all-atom explicit-solvent MD simulation under the NPT ensemble (constant number of atoms, pressure, and temperature) and 3D periodic boundary conditions. The NAMD 2.14 package (Phillips et al. 2005) and CHARMM(Brooks et al. 2009) all-atom force field (version 36m)(Huang et al. 2017) were employed in these simulations. The pressure and temperature were maintained at 1 atm and 310 K using the Langevin piston control algorithm and the Langevin thermostat method, respectively. A damping coefficient of 1 ps^-1^ was applied. All covalent bonds involving hydrogen atoms were constrained using the RATTLE method. This allowed us to employ the velocity Verlet algorithm for integrating the Newtonian motion equation with a larger time step of 2 fs. The interaction potentials between atoms were computed using the particle mesh Ewald (PME) method for long-range electrostatic interactions (grid spacing of 1.0 Å) and switching functions for short-range van der Waals (vdW) interactions (twin-range cutoff at 12 and 14 Å). The CHARMM and VMD packages(Humphrey et al. 1996) were utilized for result analysis, employing FORTRAN and TCL scripts. The root-mean-square deviation (RMSD) profiles indicated that all studied systems reached convergence after 500 ns. Averages were taken over the last 1 μs trajectories to ensure data reliability and consistency.

## ACKNOWLEDGMENTS

This project was supported in whole or in part by federal funds from the National Cancer Institute, National Institutes of Health, under contract HHSN261201500003I. The contents of this publication do not necessarily reflect the views or policies of the Department of Health and Human Services, nor does mention of trade names, commercial products, or organizations imply endorsement by the U.S. government. This research was supported [in part] by the Intramural Research Program of the NIH, National Cancer Institute, Center for Cancer Research. All simulations had been performed using the high-performance computational facilities of the Biowulf cluster at the National Institutes of Health, Bethesda, MD (https://hpc.nih.gov/).

## CONFLICT OF INTEREST

The author declares that there is no conflict of interest that could be perceived as prejudicing the impartiality of the research reported.

## Notes

### Competing Interest Statement

The authors have declared no competing interest.

