## Supplementary Material for "Cell cycle progression mechanisms: slower cyclin-D/CDK4 activation and faster cyclin-E/CDK2"

**This PDF file includes:**

Figures S1 to S5
Tables S1

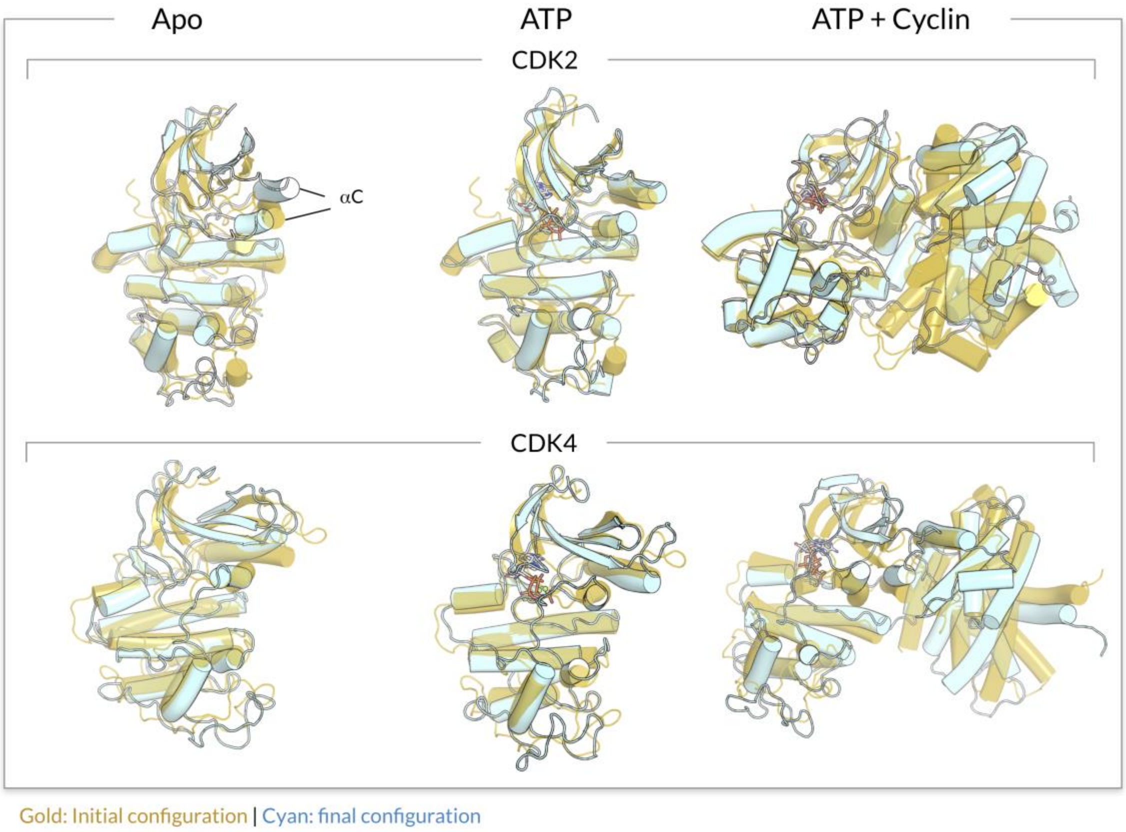


**Fig. S1.** Superimposition of the final conformation (cyan cartoon) into the starting point (gold cartoon) for the CDK2 (*upper panel*) and CDK4 (*lower panel*) systems. From left to right column, the cartoons are represented in the following order: inactive CDK2 and CDK4 monomer ("Apo"), inactive CDK2 and CDK4 monomer with ATP ("ATP"), and inactive CDK2 and CDK4 with ATP and their respective cyclin partners ("ATP + Cyclin").

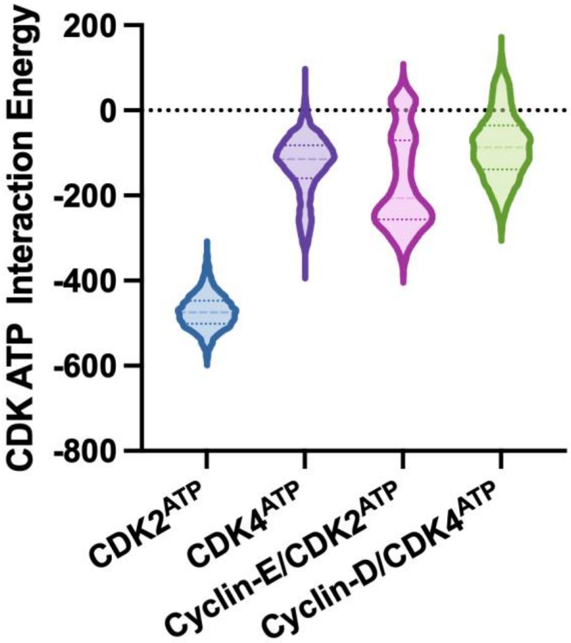


**Fig. S2.** Violin plots representing the interaction energy of ATP with the kinase domain of CDKs for the monomeric systems, $CDK2^{ATP}$ and $CDK4^{ATP}$, and the complexes, cyclin-E/$CDK2^{ATP}$ and cyclin-D/$CDK4^{ATP}$. In the monomeric CDKs, ATP exhibits a stronger interaction with $CDK2^{ATP}$ than with $CDK4^{ATP}$. For the complexes, ATP has a slightly stronger binding affinity to the CDK2 complex compared to the CDK4 complex.

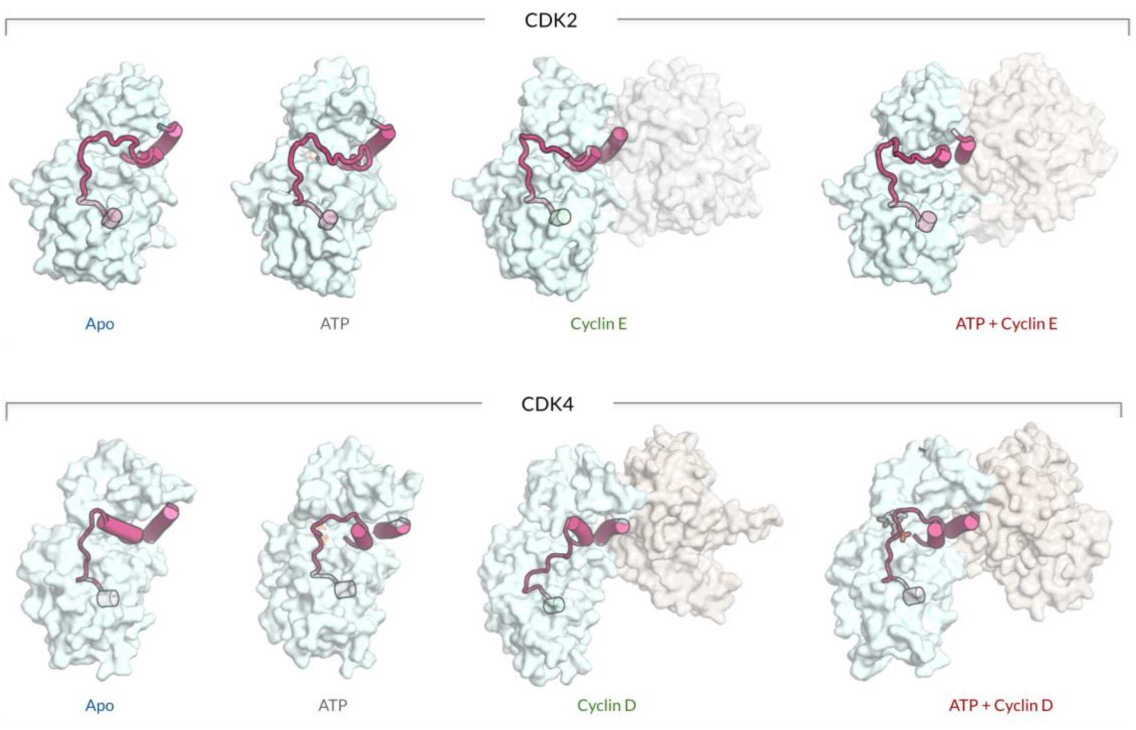


**Fig. S3**. Simulation snapshots highlighting the position of the αC-helix and A-loop. For CDK2, ATP loading or cyclin-E binding induces the outward movement of the A-loop (perpendicular to the page) and the inward movement of the αC-helix. In contrast, for CDK4, ATP loading or cyclin-D binding induces the movement of the inhibitory αL12-helix on the activation loop closer to the αC helix, while the position of the αC-helix remains relatively unchanged.

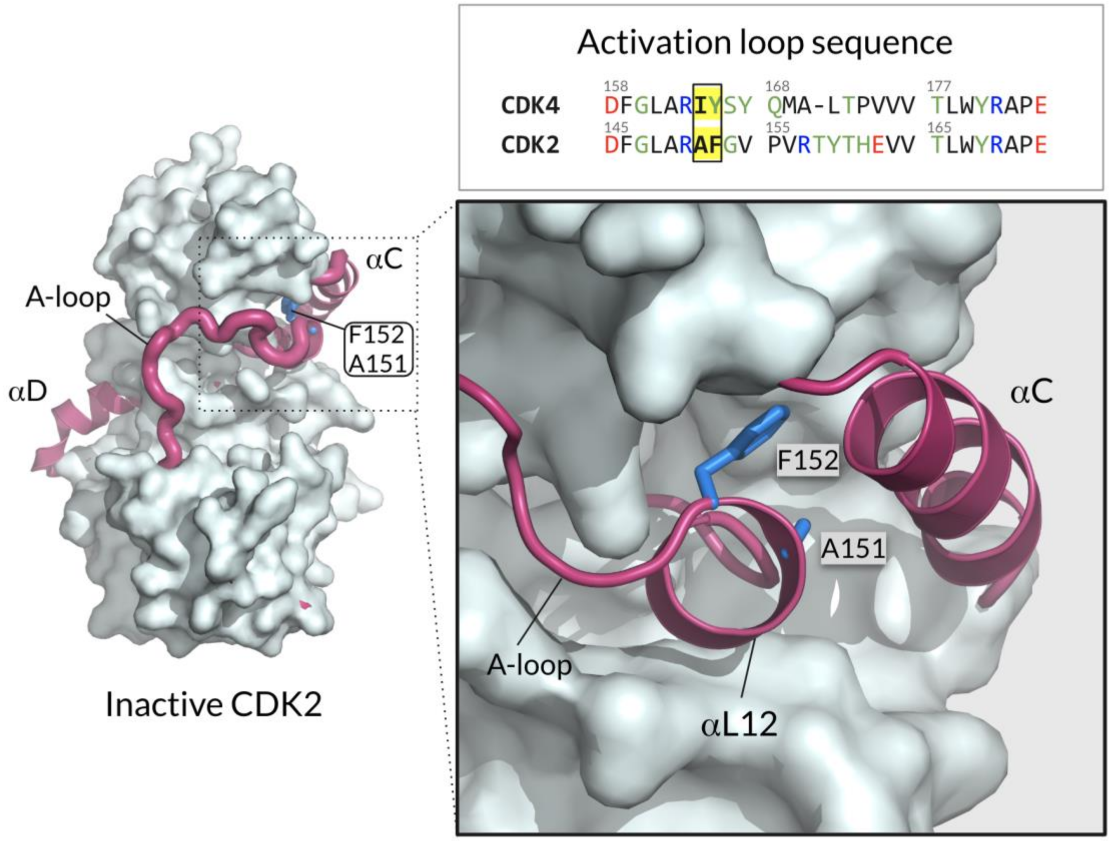


**Fig. S4.** Surface representation of the monomeric inactive CDK2$^{Apo}$ with its A-loop and αC-helix rendered as thick tube and helix, respectively (*left panel*). Highlight (*right panel*) showing the position of the αL12-helix and αC-helix. F152 and A151 on the A-loop of CDK2 are also shown.

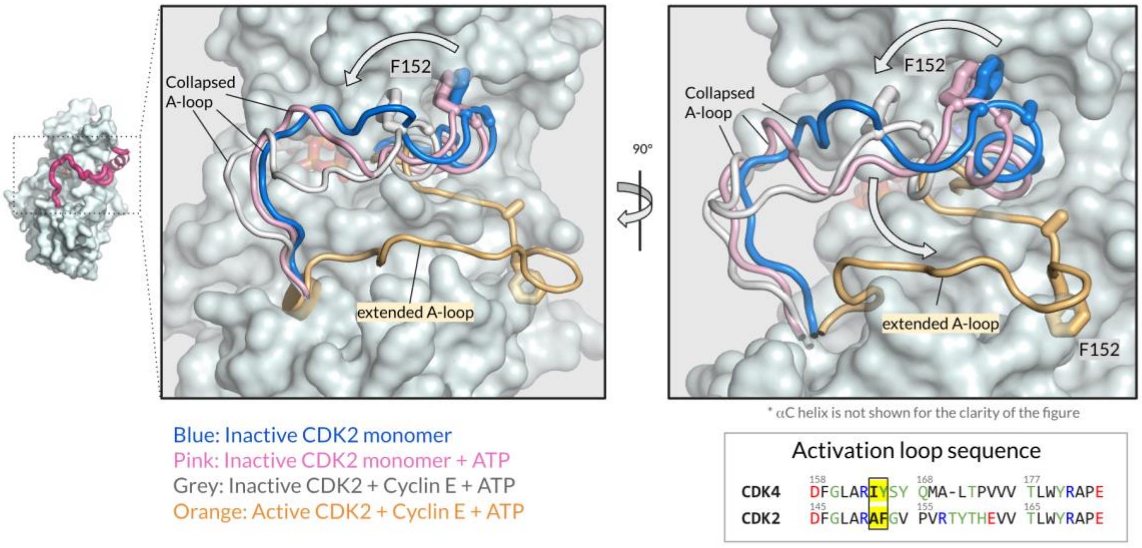


**Fig. S5.** Surface representation of the inactive CDK2$^{Apo}$ with its A-loop and αC-helix rendered as thick tube and helix, respectively (*left panel*). Superimposition of the A-loop of the CDK2 systems (*middle and right panels*). It shows the movement of the A-loop of CDK2 upon ATP loading or cyclin-E binding, or both. We observe that the A-loop of CDK2 moves outward on either ATP loading or cyclin binding.

**Table S1.** Summary of simulation systems of the CDK2 and CDK4.

| PDB | Systems | Num. of Atoms | Simulation Time (μs) |
|---|---|---|---|
| 1HCL | $CDK2^{Apo}$ | 73387 | 2 |
| 1HCK | $CDK2^{ATP}$ | 73360 | 2 |
| 1W98 + 1HCL | Cyclin-E/$CDK2^{Apo}$ | 134582 | 2 |
| 1W98 + 1HCK | Cyclin-E/$CDK2^{ATP}$ | 134635 | 2 |
| 2W9Z | $CDK4^{Apo}$ | 73360 | 2 |
| 2W9Z | $CDK4^{ATP}$ | 73413 | 2 |
| 2W9Z | Cyclin-D/$CDK4^{Apo}$ | 134407 | 2 |
| 2W9Z | Cyclin-D/$CDK4^{ATP}$ | 134355 | 2 |
